# Use of a scoring strategy to determine clinical risk of progression and risk group-specific treatment adherence in subjects with latent tuberculosis infection

**DOI:** 10.1101/207852

**Authors:** Michael Scolarici, Ken Dekitani, Ling Chen, Marcia Sokol-Anderson, Daniel F Hoft, Soumya Chatterjee

**Affiliations:** Division of Infectious Diseases, Allergy and Immunology, Department of Internal Medicine, St Louis University, St Louis, MO; St Louis University School of Medicine, St Louis, MO; Division of Biostatistics, Washington University in St. Louis School of Medicine, St Louis, MO

**Keywords:** Latent tuberculosis, risk assessment, treatment adherence

## Abstract

**Background:** Annual incidence of active tuberculosis (TB) cases has plateaued in the US from 2013-2015. Most cases are from reactivation of latent tuberculosis infection (LTBI). A likely contributor is suboptimal LTBI treatment completion rates in subjects at high risk of developing active TB. It is unknown whether these patients are adequately identified and treated under current standard of care.

**Methods:** In this study, we sought to retrospectively assess the utility of an online risk calculator (tstin3d.com) in determining probability of LTBI and defining the characteristics and treatment outcomes of Low: 0-<10%, Intermediate: 10-<50% and High: 50-100% risk groups of asymptomatic subjects with LTBI seen between 2010-2015.

**Results:** 51(41%), 46 (37%) and 28 (22%) subjects were in Low, Intermediate and High risk groups respectively. Tstin3d.com was useful in determining the probability of LTBI in tuberculin skin test positive US born subjects. Of 114 subjects with available treatment information, overall completion rate was 61% and rates of completion in Low (60%), Intermediate (63%) and High (57%) risk groups were equivalent. 75% subjects in the 3HP group completed treatment compared to 58% in the INH group. Provider documentation of important clinical risk factors was often incomplete. Logistic regression analysis showed no clear trends of treatment completion being associated with assessment of a risk factor.

**Conclusion:** These findings suggest tstin3d.com could be utilized in the US setting for risk stratification of patients with LTBI and select treatment based on risk. Current standard of care practice leads to subjects in all groups finishing treatment at equivalent rates.

## Introduction

One third of the world population is estimated to have LTBI [1], a state of infection caused by *Mycobacterium tuberculosis* (Mtb) characterized by temporary immune containment of the bacteria and lack of any clinical or microbiologic evidence of active tuberculosis (TB). As these subjects have no evidence of disease, they can currently be diagnosed only by measuring the cutaneous Delayed Type Hypersensitivity reaction (tuberculin skin test, TST) or by measuring production of Interferon-γ in the blood using interferon-γ release assays (IGRAs) [2]. It is estimated that 5-10% of all subjects with LTBI will have a lifetime risk of progression to active TB disease. Treatment of LTBI decreases the overall burden of active TB by 60-90% [3]. In low burden TB countries like the US where the overall active TB rates are <10/1000 population, the management of LTBI is a critical component of the new WHO post-2015 End TB Strategy and both the CDC and WHO recommend treatment of subjects deemed to have LTBI [4]. However, TB incidence in the US has plateaued at 3.0 cases per 100,000 persons between 2013-2015 [5].Although the reasons for this trend are not completely clear, poor rates of completion of treatment for LTBI (i.e. 50 % or less on average) [6] is a likely contributing factor. In part, poor completion rates are due to prolonged treatment required for all subjects defined to have LTBI with Isoniazid (INH) for 6-9 months or 4 months of Rifampin. Prolonged treatment with these drugs also increases chances of liver toxicity thereby limiting adherence. Recently, a 3 month regimen of weekly INH and Rifapentine (3HP) has been approved by the CDC. 3HP has shown equivalent efficacy but is resource-intensive because it requires direct observation of the patients by a healthcare worker. The CDC therefore recommends this regimen only in select clinical high risk groups [7]. Although certain clinical risk factors such as being infected with HIV, diabetes, recent contact with an active TB patient or receiving immune modulatory drugs like TNF-α blocker therapy are well known to increase the risk of developing TB disease, data are lacking on the frequency with which a comprehensive assessment of all clinical risk factors is performed in subjects with LTBI by health care providers. Provider awareness of a subject being at high risk of progression to active TB could facilitate completion of treatment in those at high risk. www.TSTin3D.com is a validated online calculator that combines TST or IGRA screening results with other clinically pertinent information, to better estimate the positive predictive value (PPV) of TB infection in a given individual. The calculator also allows for systematic assessment of additional medical risk factors to calculate an individual’s annual and cumulative risk of progression to active TB. Such assessments not only allow the identification of patients that are at high risk of progression [8] but could be used to select shorter, supervised regimens to ensure treatment completion in that group. Therefore, to assess utility of the calculator in a clinical setting, we used it to perform a retrospective systematic quantification of risk, assessed provider risk awareness and compared treatment completion rates in subjects at “Low”, “Intermediate” and “High” risk of LTBI reactivation.

## METHODS

### Study subjects and data collection

Data were collected retrospectively on 125 adult patients (≥18 years and ≤80 years) with LTBI that were seen in the Saint Louis University Infectious Diseases outpatient clinic between January 1,2010 and December 31^st^, 2015. Patients with LTBI were first screened using ICD9/10 LTBI diagnosis codes, 795.51, 795.52 and R76.11, R76.12 respectively. Patients were included only if they had a documented positive TST result available at their first visit to the clinic, were not suspected of having active TB by the clinic physician and had no prior history of treatment for LTBI or active TB. Patient electronic medical records were reviewed for variables required by the calculator (TSTin3D.com) to estimate an individual’s annual and cumulative risk of LTBI reactivation. These variables are listed in **Supplemental Methods**. Since TSTin3D.com only considers race of those born in the USA, data on race was collected only for US born individuals. Information about type and duration of antibiotic therapy and last known documented follow-up was recorded to calculate duration of treatment. For patients not able to complete the full course of LTBI therapy, information about reasons for discontinuation were obtained. The study was approved by the St Louis University Institutional Review Board. To assess how frequently healthcare providers comprehensively asses the risk factors affecting the patient’s risk of progression to active TB; we queried the history and physical documented, along with blood and radiologic testing ordered at initial visit to the Infectious Diseases clinic as noted in the electronic medical record.

### Variables generated by the calculator

Using the above data, the calculator was used to generate a positive predictive value (PPV) for TST performed to detect TB infection. As IGRAs have higher specificity for assessment of Mtb infection, the calculator assigns a PPV of 98% if IGRA is positive. The positive predictive value (PPV) of the TST is the patient’s probability of having true latent TB infection based on a positive result. Details of PPV calculations are presented in **Supplemental Methods**. TSTin3d.com also generated an annual and cumulative risks (up to age 80 years) of progression to active TB disease as well as the risk of drug induced hepatitis from INH use (details in **Supplemental Methods**). The baseline annual risk of TB disease calculated by tstin3d.com was obtained from a large cohort of healthy TST-positive US military recruits followed up for 4 years [9]. In this algorithm the highest annual risk is assigned (in descending order) to transplantation requiring immunosuppressant therapy (7.4%), HIV (5%), pulmonary silicosis (3%), chronic renal failure (requiring hemodialysis) (2.5%), carcinoma of the head and neck(1.6%), close contact of person with active TB (1.5%) and recent TST/IGRA conversion (≤ 2years) (1.5%). The cumulative risk refers to the annual risk of TB reactivation multiplied by the number of years before the patient reaches an age of 80 years. In addition, provider awareness of risk was assessed by identifying how many of the above variables were documented by the treating physician at the initial clinical encounter.

### Analysis

Concordance between TST and IGRA in subjects that received both tests was analyzed by McNemar’s test of corrected proportions. Descriptive statistics were analyzed to determine the characteristics of subjects with a PPV of greater than 50%. Patients were also stratified into Low (<10%), Intermediate (10% to <50%), and High (50%-100%) cumulative risk categories. Furthermore, percentages of patients in each cumulative risk group completing treatment were calculated. Fisher’s exact test was used to determine the association between the types of drug regimen with treatment completion rates. Logistic regression analysis was performed to examine the association between provider assessment of an individual clinical risk factor and the patient’s likelihood of completing the treatment. All statistical analyses were conducted using SAS 9.4 (SAS Institute, Cary, NC), two-sided with a significance level of 0.05.

## RESULTS

### Characteristics of the study subjects

**Table 1** shows the baseline demographic information for the study subjects. The median age was 49 years and a little less than half were female (43.31%). Of the 125 patients included, 94 were from the US and US territories, 32 were non-US born and 1 subject had no documentation of country of origin. Of the US-born individuals most were Caucasian (52%) or African American (43%). The mean age at immigration for non-US born subjects was 29.22 years. TST data were available on 69 and IGRA data on 91 subjects. Of all 35 TST positive subjects who also had IGRA results available, 19 were IGRA positive, 13 were IGRA negative and 1 indeterminate. The Only 2 subjects with positive IGRA had a negative TST result (**Figure 1**). There was significant discordance between the two tests (p = 0.0045,Mcnemar’s test) in subjects who underwent both TST and IGRA for diagnosis. There were 90 subjects who had a single test performed (either IGRA or TST) of whom 56 (44%) had a positive IGRA (either a Quantiferon-TB Gold or T-spot TB) and 34 (27%) had a positive TST. The majority of patients who received an IGRA received the QuantiFERON-TB Gold test. 5 patients received both the Quantiferon-TB Gold and the T-spot TB test.

**Figure 1.**
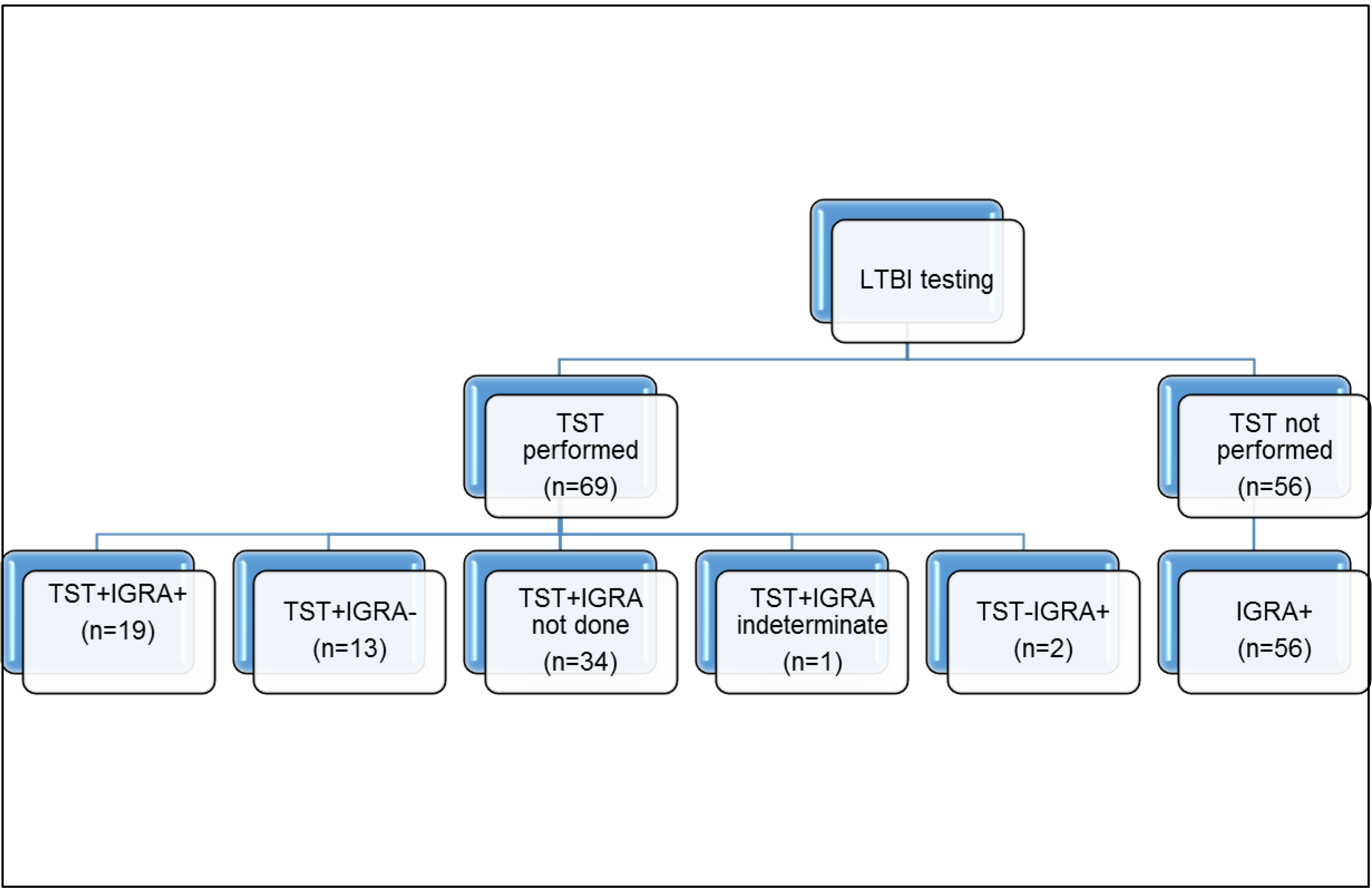
Showing all subjects included in the study based on whether they had a tuberculin skin test (TST) performed. Those who tested negative on TST or had no TST results available had to have a positive Interferon Gamma Release Assay (IGRA) i.e. Quantiferon-TB Gold or T-spot TB test to be included in the study.

**Table 1.** Characteristics of patients with latent tuberculosis infection.

| **Patient Characteristics** |  | **Number (min to max)** |
| --- | --- | --- |
| Median Age (years) |  | 49, Range:22-80 |
| Median age at immigration to USA (years) |  | 28.5 (3-45) |
| Height (cm) |  | 170.2 (147.3-195.6) |
| Weight (kg) |  | 82.5 (47.3-175.1) |
| BMI (kg/m<sup>2</sup>) |  | 29.15 (18.7-55.39) |
| **Patient demographics** |  | **Number (%)** |
| Sex (Female) |  | 54 (43.2%) |
| Ethnicity |  |  |
| Caucasian |  | 48 (38.4) |
| African American/Black |  | 39(31.2) |
| Hispanic |  | 1 |
| Other |  | 35(28) |
| Unknown |  | 2(1.6) |
| **Country of Birth** |  | **Number** |
|  | Bosnia | 4 |
|  | Burma/Myanmar | 1 |
|  | Congo | 2 |
|  | India | 3 |
|  | Iraq | 2 |
|  | Ireland | 1 |
|  | Kenya | 1 |
|  | Malaysia | 1 |
|  | Mexico | 1 |
|  | Nepal | 3 |
|  | Pakistan | 2 |
|  | Philippines | 1 |
|  | Russia | 1 |
|  | Somalia | 2 |
|  | Thailand | 1 |
|  | USA | 94 |
|  | USA-Puerto Rico | 1 |
|  | Vietnam | 3 |
|  | Unknown | 1 |
| Total |  | 125 |

### Positive Predictive Value of TST

Most foreign born subjects who underwent only TST were from high or intermediate TB burden countries and consequently had a PPV value of 50% or greater for being TST positive. Therefore, tstin3d.com was primarily useful for calculating the PPV in 22 US-born subjects who had a positive TST with either IGRA not done or indeterminate. Of the US born subjects who were diagnosed only by TST alone, higher number of Non-Hispanic Black/African-American subjects were in the >50% PPV group compared to those with PPV<50%. This was likely due to the higher risk assigned by the calculator to the Non-Hispanic Black/African-Americans with a positive TST.

### Risk of Reactivation

The overall distribution of risk factors in the study patients comparing US born with immigrants is shown in **Figure 2**. Subjects with HIV, diabetes, history of smoking, taking a TNF-α inhibitor and on transplant immunosuppression were more common in US born subjects compared to immigrants. The calculator allowed us to divide subjects with LTBI into “Low (<10% risk),” “Intermediate (10-<50% risk),” and “High (50-100% risk)” cumulative risk groups. All patients with AIDS had the highest annual (>10%) and cumulative (>50%) risk. The calculator assigns a slightly increased risk to African Americans compared to age and sex matched Caucasians and a larger proportion of African Americans (20% and 43%) were in the highest annual (>10%) and cumulative risk category (>50%) compared to Caucasians (2% and 16%). However, this was primarily because of the significantly increased prevalence of HIV/AIDS noted in this population (15/42, 37.5%) compared to the Caucasian population (4/49, 8.2%) (p=0.001 Fisher exact test). **Table 2** shows characteristics of all the patients in the High cumulative risk group. Furthermore, for all patients who tested positive on TST but negative on IGRA, the median cumulative risk was 19% suggesting that they were not in the high risk group.

**Figure 2.**
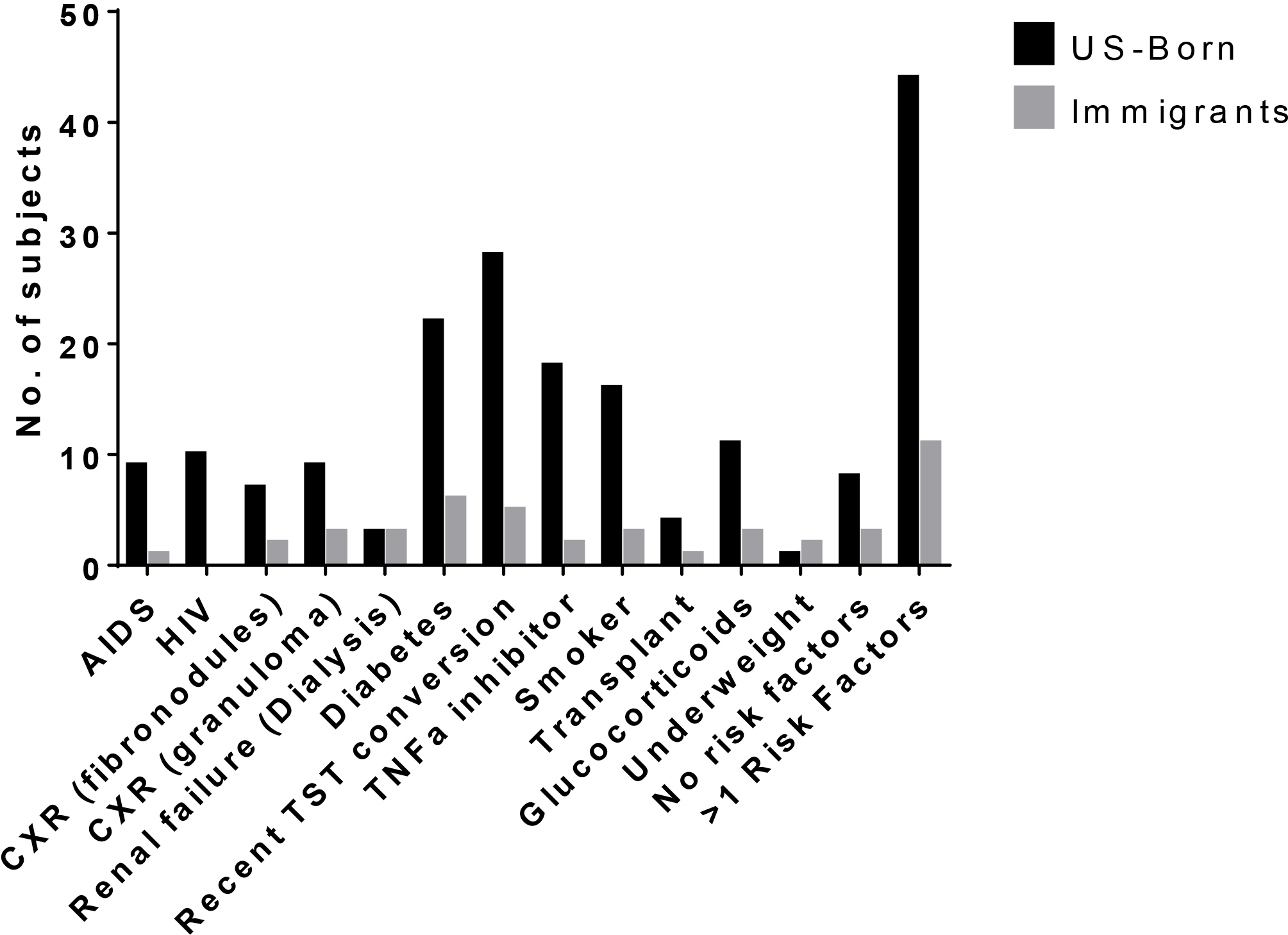
Showing number of US born (black bars) vs. non-US born (grey bars) subjects with different risk factors for progression to active TB disease in the overall cohort of patients with latent TB

**Table 2.** Distribution of medical risk factors for progression to active TB in subjects in the High (50-100% cumulative risk) group.

| **Risk factor** | **Number** | **Cumulative risk value%** |
| --- | --- | --- |
| **HIV (non AIDS)** | 10 | 100 (Median risk)* |
| • No comorbidities<br>• Smoker<br>• Recent TST conversion ($\leq 2$ years ago) | 0<br>2<br>9 |  |
| **AIDS** | 10 | 100 (Median risk)* |
| • No comorbidities<br>• Fibronodular disease on chest X-ray +Recent TST conversion ($\leq 2$ years ago)+renal failure on dialysis<br>• Smoker<br>• Smoker+ Recent TST conversion ($\leq 2$ years ago)<br>• Recent TST conversion ($\leq 2$ years ago)<br>• Diabetes | 2<br>1<br>4<br>2<br>6<br>1 |  |
| Recent TST conversion ($\leq 2$ years ago)+renal failure on dialysis | 1 | 85.04 |
| Renal failure on dialysis | 1 | 70.3 |
| Recent TST conversion + TNF-α blocker + steroid use | 2 | 51.69, 8.61 |
| Immunosuppression after solid organ transplant | 2 | 100, 87.51 |
| Diabetes+ steroid use + solid organ transplant | 1 | 100 |
| Fibronodular disease on chest X-ray + solid organ transplant | 1 | 69.97 |
| Fibronodular disease on chest X-ray + bone marrow transplant | 1 | 76.15 |

**\*All subjects with HIV and/or AIDS had 100% cumulative risk**

### Risk Factors and Treatment Completion in Specific High risk groups Subjects on TNF-α blockers and LTBI

Data were available for 20 subjects with LTBI on TNFα blockers. A significantly higher proportion (75%) of subjects on TNFα blockers (TNFα group) were seen because of recent (< 2 years ago) TST/IGRA conversion from negative to positive, compared to 17.5% in the control group (76 HIV-negative subjects not on any immunosuppressive therapy, p=0.002, Fisher exact test). The distribution of other pertinent risk factors were as follows between the TNFα group and control group: diabetes (6% vs. 26%), smoking >1 pack per day (6% vs. 11%), and renal failure (0% vs. 6%). The median cumulative risk of progression to active TB was 17.25% in the TNFα group (range: 8.6 – 51.7%) and 5.6% in the control group (range: 0.1 – 100%). 70% of patients were able to successfully complete LTBI therapy in the TNFα group. This could be due to close follow up in the rheumatology clinic.

### Subjects with HIV and LTBI

Of the HIV positive population with LTBI, 10 of 20 had AIDS. The median annual TB-risk amongst HIV, AIDS, and HIV-uninfected patients was 8% (3-8%), 22% (21-25%), and 0.5% (0-6%). More patients with HIV/AIDS smoked >1 pack-per-day (HIV/AIDS: 30%, HIV-uninfected: 13%). Other risk-factors distributed amongst HIV, AIDS and HIV-uninfected groups: recent TST/IGRA conversion (10%, 30%, and 28%), renal-failure requiring hemodialysis (0%, 10%, and 0%), and diabetes (0%, 10%, 25 %). Most HIV positive patients were prescribed 9H (HIV/AIDS: 85%, HIV-uninfected: 39 %). The median treatment-days were longer (274 vs. 117 days) but the rates of treatment completion were comparable for HIV/AIDS than HIV-uninfected patients (30% vs. 34%). Reasons HIV/AIDS patients stopped therapy included loss to follow-up (2/6), cost (1/6), hepatitis (1/6), rash (1/6), and clinical contraindications (1/6).

### Hepatitis while on LTBI treatment

The calculated risk of INH induced hepatitis correlates with the age of the patient more than any of the other risk factors. Of the five patients that developed elevated LFTs, all were above 38 years of age with three being above 65 years. Only one was on INH and four were on rifampin. The calculator estimated a risk of hepatitis of 1.2% for two of these subjects, 5% for two and 2.3% for one subject. Each patient was concomitantly using at least one other potentially hepatotoxic medication.

### Treatment completion

Of the 125 subjects included in the analysis, 8 were not offered any treatment and 3 refused. 59 of 114 (52%) subjects were started on treatment with INH, 24 (21%) on 3HP, 23 (20%) on Rifampin and 8 (7%) on Rifabutin. 69 (61%) subjects completed the recommended duration of therapy during the study period and none developed active TB. The completion rates of those treated for LTBI were the best for 3HP and worst for Rifabutin, 75% and 50% subjects respectively (**Table 3**). Amongst the documented reasons for stopping treatment early (**Figure 3**), loss to follow-up accounted for a majority of incomplete treatments (23 subjects). Other reasons for stopping that were categorized were: elevated liver enzymes, cost/access, minor side effects (nausea, vomiting, and diarrhea), rash, drug interactions and clinical decision to stop. No reasons for stopping were documented in 4 subjects. Importantly, no significant differences were observed between patients completing therapy therapy in the Low, Intermediate and High cumulative risk groups as shown in **Table 4** (p=0.54,Chi-square test for trends).

**Figure 3.**
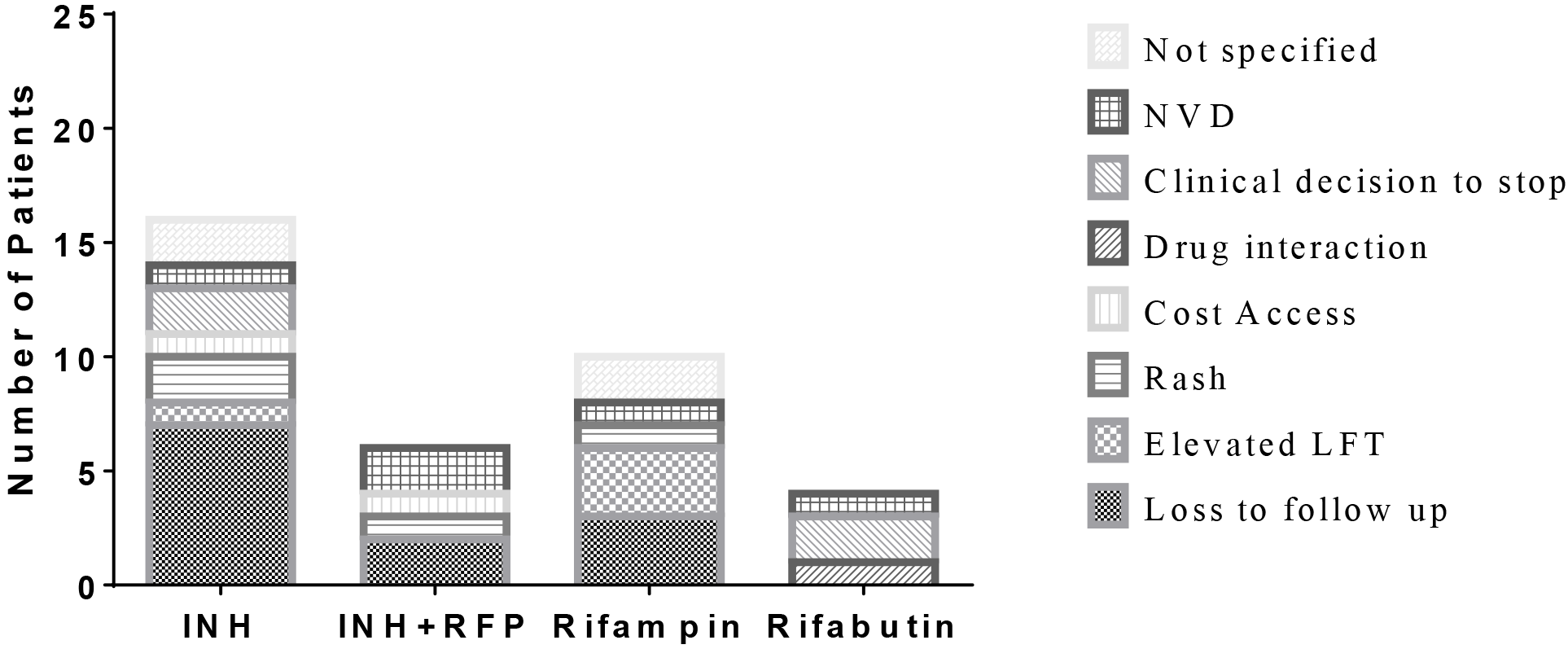
Showing number of patients discontinuing therapy and the primary documented causes by drug group.

**Table 3.** Subjects completing treatment by drug category.

| **Drug Regimen** | **Total number, n** | **Completed treatment, n (%)** |
| --- | --- | --- |
| INH | 59 | 34 (58) |
| 3HP | 24 | 18 (75) |
| Rifampin | 23 | 13 (57) |
| Rifabutin | 8 | 4 (50) |

**Table 4.** Subjects completing treatment for latent tuberculosis divided by risk group.

| **Cumulative risk group (RG)** | **Total number, n** | **Number started on treatment, n** | **Completed Treatment, n (% of those started)** |
| --- | --- | --- | --- |
| Low (0 - <10%) | 51 | 45 | 27 (60) |
| Intermediate (10 - <50%) | 46 | 41 | 26 (63) |
| High (50 - 100%) | 28 | 28 | 16 (57) |

### Provider risk documentation

For analysis of whether a specific risk factor was assessed by the ID clinic provider, we queried the electronic medical record for documentation of risk factors in the written clinical assessment by the health care provider. As shown in **Table 5**, BCG status was documented for only 15 of 32 immigrants. Less than half of the patients’ initial encounter included documented information about country of birth, HIV/AIDS status, recent TST/IGRA conversion, young age at infection, recent TST/IGRA conversion (≤2 years ago) and history of cancer. Hyperglycemia and renal failure were assumed to be assessed if the provider had access to a blood test for basic metabolic panel at or within 3 months prior to seeing the patient. Most patients were assessed for smoking and malnutrition and had received a chest X-ray at or within 3 months of their Infectious Diseases clinic visit. As shown in **Table 6**, logistic regression analysis showed that there was no statistically significant association of risk factor assessment in the clinic at initial visit with the probability of subjects completing treatment. However, as shown by the higher Odds Ratios, documentation of immunosuppression (HIV/AIDS, steroids, transplant related immunosuppression) as well as documentation of smoking and recent (≤ 2 years) TST/IGRA conversion were associated with a better chance of completing treatment.

**Table 5.**
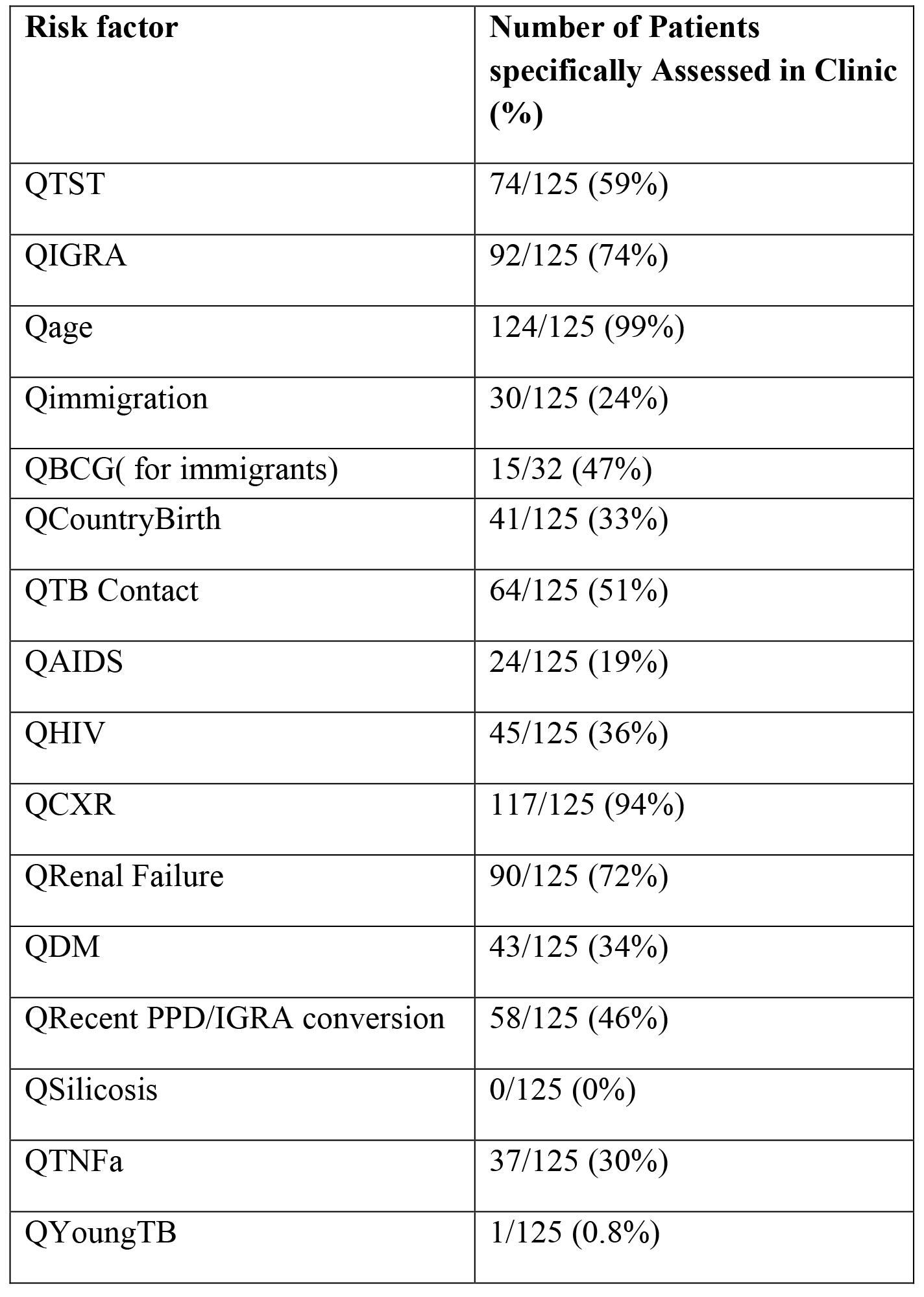
Subjects with latent tuberculosis who were assessed for the different risk factors by Infectious Diseases clinic providers.

| **Risk factor** | **Number of Patients specifically Assessed in Clinic (%)** |
| --- | --- |
| QTST | 74/125 (59%) |
| QIGRA | 92/125 (74%) |
| Qage | 124/125 (99%) |
| Qimmigration | 30/125 (24%) |
| QBCG( for immigrants) | 15/32 (47%) |
| QCountryBirth | 41/125 (33%) |
| QTB Contact | 64/125 (51%) |
| QAIDS | 24/125 (19%) |
| QHIV | 45/125 (36%) |
| QCXR | 117/125 (94%) |
| QRenal Failure | 90/125 (72%) |
| QDM | 43/125 (34%) |
| QRecent PPD/IGRA conversion | 58/125 (46%) |
| QSilicosis | 0/125 (0%) |
| QTNFa | 37/125 (30%) |
| QYoungTB | 1/125 (0.8%) |

**Table 6.** Logistic regression analysis of relating probability of completion of treatment to assessment of selected individual TB progression risk factors.

| **Assessed Risk factor\*** | **Odds Ratio (Confidence Interval)** | **p value** |
| --- | --- | --- |
| Renal failure | 0.83 (0.35-2.0) | 0.67 |
| HIV | 1.83 (0.83-4) | 0.13 |
| AIDS | 1.36 (0.53-3.5) | 0.52 |
| Chest X-ray available | 0.36 (0.082-1.6) | 0.18 |
| Diabetes mellitus | 0.89 (0.4-1.9) | 0.75 |
| Recent TST conversion ($\leq 2$ years ago) | 0.78 (0.37-1.7) | 0.51 |
| On TNF-α blocker | 0.72 (0.31-1.7) | 0.45 |
| Smoker | 1.2 (0.43-3.4) | 0.72 |
| On steroids | 0.99 (0.39-2.5) | 0.97 |
| Transplant immunosuppression | 1.56 (0.2-11.4) | 0.67 |

\*Young age at TB and Head and Neck cancer are not included as number of subjects assessed were not enough to include in logistic regression analysis

## Discussion

In this study we retrospectively applied the validated risk calculator (tstin3d.com), to describe the distribution of clinical risk factors contributing to increased annual and cumulative risk of progression to active TB disease, in patients with LTBI seen at the Saint Louis University outpatient Infectious Diseases (ID) clinic over a 5 year time period. We were also able to measure rates of treatment completion in the groups with differing cumulative risk. Finally, we measured the frequency with which different risk factors were documented at initial visit by the healthcare provider in the ID clinic. Subjects were included only if they had a documented positive TST/IGRA result available at or prior to their clinic visit, had no clinical evidence of active TB (as documented by the ID clinic physician) and had no prior history of treatment for either active TB or LTBI. Our study demonstrates that tstin3d.com can be used for systematic assessment of major risk factors for progression to active TB in subjects with LTBI and divide subjects with LTBI into Low, Intermediate and High cumulative risk groups based on the presence or absence of a combination of clinical risk factors. Our data show that current standard of care practice did not result in higher treatment completion rates for patients in the High risk group-an important goal of TB control in low burden countries. Evaluation of any particular risk factor was not associated with improved treatment completion but this could be because providers often select regimens based on side effect profiles of drugs rather than the patient’s risk of progression to active TB. We speculate that provider awareness of a numerical risk for patients with LTBI can allow them to use short course LTBI treatment regimens more cost-effectively, ensuring completion in the highest risk group, an approach that needs to be tested in future prospective studies.

For patients undergoing TST, the calculator (tstin3d.com) facilitates interpretation of risk by taking into account the TST size, PPV of TST and risk of development of disease. A PPV >50% leads to an increased cumulative risk. A default PPV>98% is assigned, however, by the calculator in those with a positive IGRA. This is one limitation of the calculator as recent studies have shown that false positive IGRAs remain a concern among specific groups assessed for LTBI [10].We found significant discordance between PPD and IGRA positivity in our study in keeping with results obtained by others [11, 12]. We found the PPV calculated by the calculator to be most useful in stratifying US born subjects who underwent TST as their sole test for assessment of TB infection as BCG vaccination is not routinely offered in the US. Although the calculator was primarily developed to overcome the limitations of TST interpretation [8], we utilized it firstly, to retrospectively assess the percentage of patients with LTBI in Low, Intermediate and High individual cumulative risk groups. We then measured the percentage of patients finishing LTBI treatment in the different risk groups. We noted equivalent rates of treatment completion between the different risk groups which raises the possibility thattstin3d.com use by providers might help select targeted strategies to ensure completion in the high risk group. ID clinic assessment of any particular risk factor was not associated with treatment completion but this might be due to the relatively small sample size of our population. However, we noted non-significant but higher odds of completing treatment in patients assessed for immunosuppression and recent TST/IGRA conversion. Current CDC guidelines define specific risk groups which should be given high priority for LTBI treatment based on TST size and/or IGRA positivity and clinical risk factors [13]. However, as shown in our study, most of these risk factors may not be systematically assessed by the busy clinician in all patients with LTBI. Freely available web based tools like tstin3d.com may therefore, facilitate a comprehensive assessment of risk factors for TB progression [6]. This, in turn, may affect the choice of regimen and the intensity of follow up provided to the patients when healthcare providers become aware of those at high cumulative risk of progression. Although lack of adequate healthcare as well as patient related factors have been reported to affect adherence to LTBI treatment [6, 14, 15], specific interventions like shorter duration of therapy using Rifamycin based regimens and directly observed treatment (DOT) are well recognized measures to improve treatment adherence [16–18]. This approach also minimizes the loss to follow up which was the major reason for incomplete treatment in our cohort.

In agreement with the published literature on adherence with 3HP based regimens, we found the highest rates of treatment completion in subjects prescribed 3HP compared to those receiving INH or Rifampin. Surprisingly, we saw low rates of treatment completion seen with Rifampin or Rifabutin only regimens. Cytochrome P (CYP) 450 isoenzyme induction by Rifamycin based regimens remains a concern in subjects with HIV on ARVs. This is especially important as more than 85% of subjects with HIV/AIDS were prescribed INH and only 30% were able to complete treatment in our study. Current US guidelines recommend use of 3HP only with efavirenz (EFV) - or raltegravir (RAL)-based regimens (in combination with either abacavir/lamivudine [ABC/3TC] or tenofovir disoproxil fumarate/emtricitabine [TDF/FTC]) for treatment of LTBI [18]. Allowing for drug resistance testing and patient adherence factors, a future strategy of temporarily switching HIV positive patients to the above mentioned regimens could allow better treatment completion rates with a 3HP based regimen.

Subjects on TNF-α blockers are another high risk group at risk for TB disease reactivation. The majority of subjects with LTBI in our study were referred because of recent TST/IGRA conversion (within ≤ 2years) which put them at an even higher risk of TB reactivation disease. Rifamycin based regimens have recently been shown to be effective with minimal side effects in this group of patients [19]. Although only 25% of patients on TNF-α blockers were prescribed 3HP, 70% were able to successfully complete LTBI therapy. This is likely due to close clinic follow-up that these patients receive for their underlying autoimmune disease.

Two important limitations of our study are the retrospective nature of the study and that subjects already receiving care for co-morbid conditions at different clinics were referred to our Infectious Diseases clinic, making them more likely to seek healthcare. These subjects are therefore more likely to have higher rates of LTBI treatment completion compared to the overall population of subjects with LTBI. Furthermore, as patients were often referred from community and other specialty clinics, ID clinic providers were often aware of medical comorbidities for e.g. HIV/AIDS, Diabetes, renal failure on dialysis, and ongoing use of TNF-α blocker use at initial clinic assessment. Therefore we could not properly assess the actual frequency with which these risk factors would be assessed had the provider not been made aware beforehand. Nevertheless, we used relatively strict criteria for inclusion of subjects with previously untreated LTBI in our study and had follow up data on the majority of our study cohort. Our study suggests that tstin3d.com could be used in future prospective studies for determining a numerical risk of TB progression in patients with LTBI for improved provider awareness of those at high risk. Furthermore, prospective design would also allow for testing whether treatment completion rates can be improved by “risk score targeted” treatment (i.e. selecting a 3HP based regimen for all subjects at high cumulative risk). The utility of “risk score targeted” treatment as a strategy for decreasing the community burden of TB in the US needs to be validated in future prospective studies.

## Acknowledgements

The authors would like to thank Dr. Madhukar Pai and Dr. Dick Menzies for allowing open use of tstin3d.com. This research was supported in part by a HIV Medical Students Program award awarded to MS by the HIV Medicine Association of the Infectious Diseases Society of America.

## References

1. Corbett, E.L., et al., The Growing Burden of Tuberculosis. Archives of Internal Medicine, 2003. 163(9): p. 1009.

2. Campion, E.W., et al., LatentMycobacterium tuberculosisInfection. New England Journal of Medicine, 2015. 372(22): p. 2127–2135.

3. Lobue, P. and D. Menzies, Treatment of latent tuberculosis infection: An update. Respirology, 2010. 15(4): p. 603–22.

4. Getahun, H., et al., Management of latent Mycobacterium tuberculosis infection: WHO guidelines for low tuberculosis burden countries. The European respiratory journal, 2015. 46(6): p. 1563–1576.

5. Salinas, J.L., et al., Leveling of Tuberculosis Incidence - United States, 2013-2015. MMWR Morb Mortal Wkly Rep, 2016. 65(11): p. 273–8.

6. Li, J., et al., Adherence to treatment of latent tuberculosis infection in a clinical population in New York City. International Journal of Infectious Diseases, 2010. 14(4): p. e292–e297.

7. Recommendations for use of an isoniazid-rifapentine regimen with direct observation to treat latent Mycobacterium tuberculosis infection. MMWR Morb Mortal Wkly Rep, 2011. 60(48): p. 1650–3.

8. Menzies, D., et al., Thinking in three dimensions: a web-based algorithm to aid the interpretation of tuberculin skin test results. Int J Tuberc Lung Dis, 2008. 12(5): p. 498–505.

9. Comstock, G.W., L.B. Edwards, and V.T. Livesay, Tuberculosis morbidity in the U.S. Navy: its distribution and decline. Am Rev Respir Dis, 1974. 110(5): p. 572–80.

10. Moses, M.W., et al., Serial testing for latent tuberculosis using QuantiFERON-TB Gold In-Tube: A Markov model. Sci Rep, 2016. 6: p. 30781.

11. Talati, N.J., et al., Poor concordance between interferon-γ release assays and tuberculin skin tests in diagnosis of latent tuberculosis infection among HIV-infected individuals. BMC Infectious Diseases, 2009. 9: p. 15–15.

12. Ribeiro-Rodrigues, R., et al., Discordance of Tuberculin Skin Test and Interferon Gamma Release Assay in Recently Exposed Household Contacts of Pulmonary TB Cases in Brazil. PLoS ONE, 2014. 9(5): p. e96564.

13. Targeted tuberculin testing and treatment of latent tuberculosis infection. American Thoracic Society. MMWR Recomm Rep, 2000. 49(RR-6): p. 1–51.

14. Makanjuola, T., H.B. Taddese, and A. Booth, Factors Associated with Adherence to Treatment with Isoniazid for the Prevention of Tuberculosis amongst People Living with HIV/AIDS: A Systematic Review of Qualitative Data. PLoS ONE, 2014. 9(2): p. e87166.

15. Malejczyk, K., et al., Factors associated with noncompletion of latent tuberculosis infection treatment in an inner-city population in Edmonton, Alberta. The Canadian Journal of Infectious Diseases & Medical Microbiology, 2014. 25(5): p. 281–284.

16. Sterling, T.R., et al., Three months of weekly rifapentine and isoniazid for treatment of Mycobacterium tuberculosis infection in HIV-coinfected persons. Aids, 2016. 30(10): p. 1607–15.

17. Sterling, T.R., et al., Three months of rifapentine and isoniazid for latent tuberculosis infection. N Engl J Med, 2011. 365(23): p. 2155–66.

18. Stuurman, A.L., et al., Interventions for improving adherence to treatment for latent tuberculosis infection: a systematic review. BMC Infectious Diseases, 2016. 16: p. 257.

